# Brief Peppermint Odour Exposure and Immediate Free Recall of a Visual Object Array: A Three-Arm Randomised Experiment

**DOI:** 10.64898/2026.10.31.763290

**Authors:** Barış Akar, Tuğba Altıntaş

**Author notes:** Corresponding author: Barış Akar.

## Abstract

**Background:** Odours reach limbic and prefrontal targets through a uniquely direct pathway and are often claimed to enhance cognition, yet experimental evidence for specific essential oils is mixed and frequently confounded by expectancy. We tested whether a brief inhalation of peppermint or clove essential oil, compared with no odour, alters immediate free recall of a visual object array.

**Methods:** Sixty healthy adults aged 18–50 years were randomly allocated to peppermint oil, clove oil or no odour (*n* = 20 each). After 30 s of inhalation (or none), participants studied a 4 × 4 array of 16 pictured items for 20 s and then wrote down every item they could recall, without time limit. Groups were compared with one-way and Welch ANOVA, Welch-corrected pairwise tests with Holm correction, Hedges’ *g* with 95% confidence intervals and Bayes factors.

**Results:** Recall differed across conditions (Welch *F*(2, 37.0) = 8.53, *p* < .001; ω^2^ = 0.14). Participants exposed to peppermint recalled more items (11.20 ± 2.14) than those exposed to no odour (8.45 ± 2.04; *g* = 1.29, 95% CI [0.61, 1.97], *p*(Holm) < .001, BF^10^ = 134). Clove (9.55 ± 3.19) did not differ reliably from either no odour (*g* = 0.40, *p*(Holm) = .203) or peppermint (*g* = 0.60, *p*(Holm) = .127; BF^10^ = 1.29).

**Conclusions:** A single brief exposure to peppermint oil was associated with substantially better immediate recall than no odour. Because the peppermint–clove contrast was inconclusive and no placebo odour was used, the data do not establish an odour-specific effect; general arousal and expectancy remain plausible contributors. Pre-registered, laboratory-based replications with placebo odours, blinding and arousal measures are required.

## 1. Introduction

Olfaction is unusual among the senses in its anatomy. Odour information reaches the piriform cortex, amygdala and entorhinal cortex with few intervening synapses and without an obligatory thalamic relay, giving odours privileged access to circuits that support emotion and memory^1–3^. Even the rhythm of nasal breathing entrains limbic oscillations and modulates memory performance^4^. Consistent with this anatomy, odour-evoked autobiographical memories are unusually emotional and vivid^5–7^, and they engage the amygdala and hippocampus more strongly than equivalent visual cues^8^. Odours can also act as contextual cues that improve retrieval when reinstated at test^9^.

A distinct question is whether odours can enhance cognition in the moment—that is, whether a brief exposure changes how efficiently new information is encoded. Several controlled studies suggest that some essential oils have acute, odour-specific effects. Rosemary aroma enhanced memory quality and alertness whereas lavender reduced working memory performance^10^, and plasma levels of the rosemary terpene 1,8-cineole correlated with cognitive performance after exposure^11^. Peppermint aroma improved memory and alertness relative to ylang-ylang and no odour^12^, and in a placebo-controlled study peppermint essential oil improved performance on demanding cognitive tasks and reduced mental fatigue^13^. Essential oils can also modulate reaction-time measures of alertness^14^, and long-term olfactory enrichment has been associated with improved memory and changes in the uncinate fasciculus in older adults^15^. Clove odour has received less attention; early work found effects of lavender and clove odours on mood and cognition that were modest and task-dependent^16^.

Two mechanisms are commonly proposed for such acute effects. First, peppermint contains menthol, which stimulates trigeminal cooling receptors in the nasal mucosa as well as the olfactory system^17,18^. Trigeminal and olfactory stimulation may increase arousal, and moderate increases in arousal—through locus coeruleus–noradrenergic gain modulation—can enhance attention and encoding^19,20^. Second, odours carry expectations: people believe that peppermint is invigorating and that some scents are relaxing, and such beliefs can produce effects that are independent of the odour itself^21–23^. Ambient odours can likewise change mood and anxiety in naturalistic settings^24^. Distinguishing pharmacological, arousal-related and expectancy-related routes requires appropriate control conditions.

We conducted a three-arm randomised experiment to test whether a brief inhalation of peppermint or clove essential oil, compared with no odour, alters immediate free recall of a visual array of pictured items. Peppermint was chosen as a stimulating, trigeminally active odour with prior evidence of cognitive benefits^12,13^, and clove as a familiar, non-mentholated comparison odour. Recall of a 16-item array exceeds the three-to-four-item limit of short-term storage^25^ and draws on rapid encoding into longer-lasting visual memory, which has a very large capacity for object information^26,27^; it is therefore sensitive to the efficiency of encoding during study.

## 2. Materials and Methods

### 2.1. Participants

Sixty healthy volunteers aged 18–50 years took part. Inclusion criteria were adequate literacy and normal or corrected-to-normal vision, and participants reported no diagnosis of neurological, psychiatric or cognitive disorder. Participants were not selected on the basis of sex, education or olfactory sensitivity. They were randomly allocated in equal numbers to the peppermint, clove or no-odour condition (*n* = 20 each). With 20 participants per group, the study had 80% power (α = .05) to detect pairwise differences of *d* ≥ 0.91 and an omnibus effect of Cohen’s *f* ≥ 0.41; smaller effects could not be detected reliably^28^. All participants were informed about the procedure and confidentiality and gave written informed consent. The study was approved by the Non-Interventional Research Ethics Committee of Üsküdar University (approval documentation is held by the authors and available on request) and conducted in accordance with the Declaration of Helsinki. Reporting follows the CONSORT 2010 guidance for parallel-group randomised studies where applicable^29^.

### 2.2. Odour stimuli

Two commercially available essential oils were used: peppermint (*Mentha × piperita*) and clove (*Syzygium aromaticum*). Oils were kept in their original bottles. Participants in the odour conditions inhaled directly from the open bottle held approximately 5 cm from the nose for 30 s. Participants in the no-odour condition did not receive an olfactory stimulus or a placebo bottle.

### 2.3. Memory task

The study material was a coloured 4 × 4 array containing 16 items: 14 pictured objects from varied categories (animals, fruits, household items, vehicles and a musical instrument), one letter and one digit (Figure 1A). The same array, with identical size and item order, was used for all participants. Immediately after odour exposure (or at the corresponding point in the no-odour condition), the array was displayed for 20 s and participants were asked to study it carefully. The array was then removed and participants wrote down all items they could recall on a blank A4 sheet, in any order and without time limit (Figure 1B). Each correctly named item scored one point (range 0–16).

**Figure 1.**
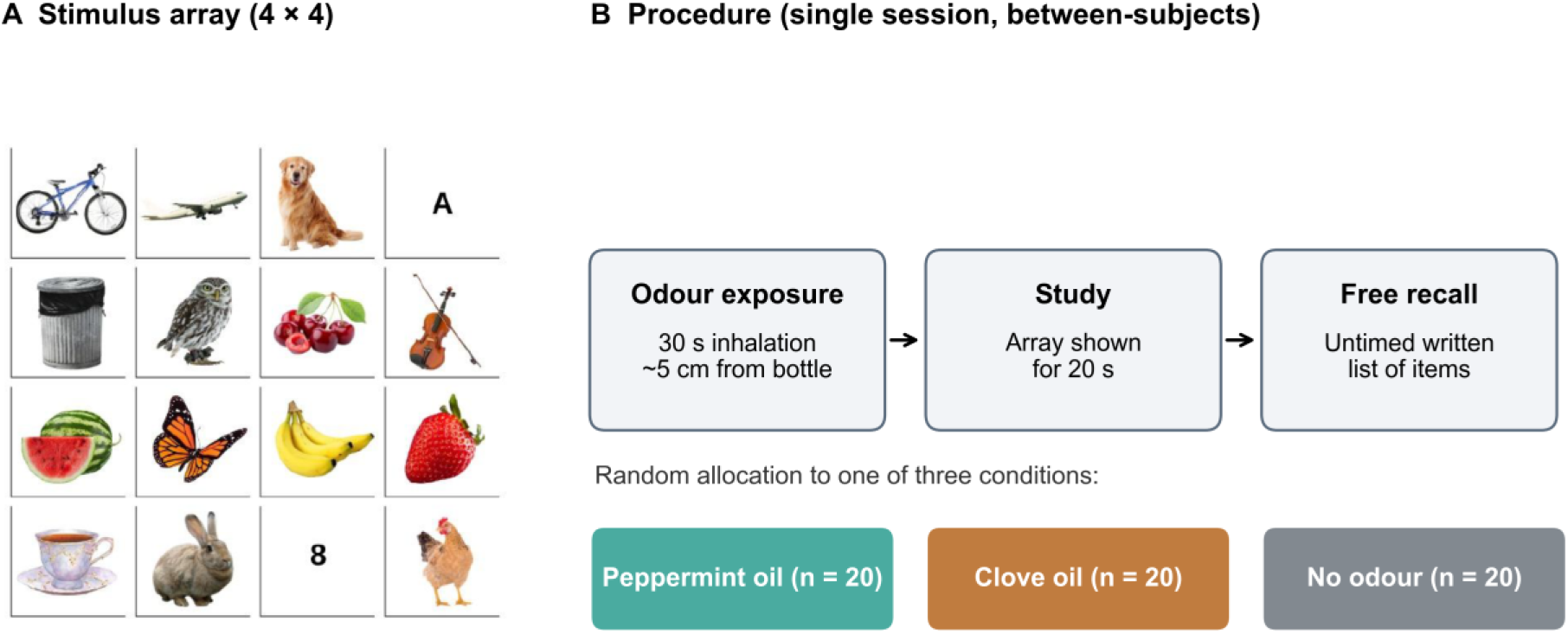
Stimulus and procedure. (A) The 4 × 4 study array (14 pictured objects, one letter and one digit) presented for 20 s. (B) Session structure. Participants were randomly allocated to a 30-s inhalation of peppermint or clove essential oil or to no odour, studied the array and then performed untimed written free recall.

### 2.4. Setting

Sessions were conducted individually in a quiet room of each participant’s home, free from distraction, with the experimenter present. Lighting, viewing distance and ambient odour could therefore not be standardised to laboratory level (see Limitations).

### 2.5. Statistical analysis

Because participant-level data were not available for this re-analysis, all statistics were computed from the group means, standard deviations and sample sizes. The effect of condition was tested with a one-way analysis of variance and, because standard deviations differed between groups (largest/smallest ratio = 1.56), with Welch’s heteroscedasticity-robust ANOVA^30^. Effect sizes are reported as η^2^ and the less biased ω^231^. Pairwise comparisons used Welch’s *t* tests^32^ with Holm correction for the three comparisons; mean differences are reported with 95% confidence intervals and standardised differences as Hedges’ *g* with 95% confidence intervals^31^. Default JZS Bayes factors (BF^10^) quantified the relative evidence for a difference versus no difference, with BF^10^ < 1/3 and > 3 taken as moderate evidence for the null and the alternative, respectively^33^. α = .05 (two-tailed). Analyses were run in Python 3.10 (SciPy, statsmodels, pingouin).

## 3. Results

Descriptive statistics are shown in Table 1 and Figure 2A. Recall differed across conditions (one-way ANOVA: *F*(2, 57) = 6.08, *p* = .004, η^2^ = 0.18, ω^2^ = 0.14; Welch ANOVA: *F*(2, 37.0) = 8.53, *p* < .001).

**Table 1.**
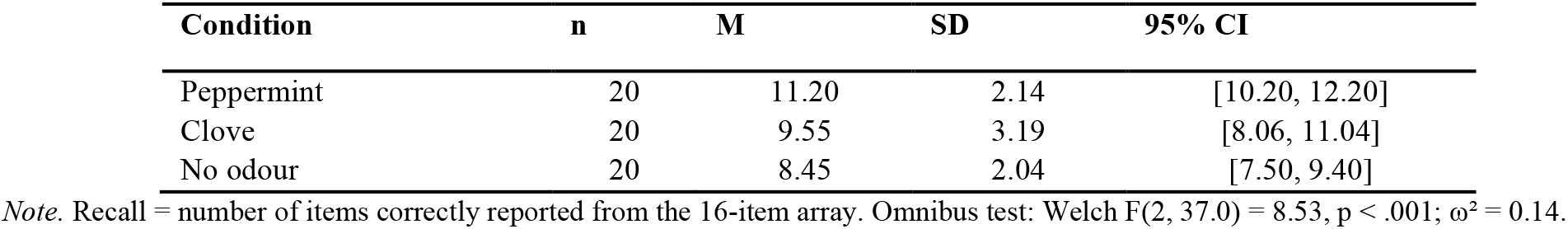
Immediate Free Recall by Condition.

**Figure 2.**
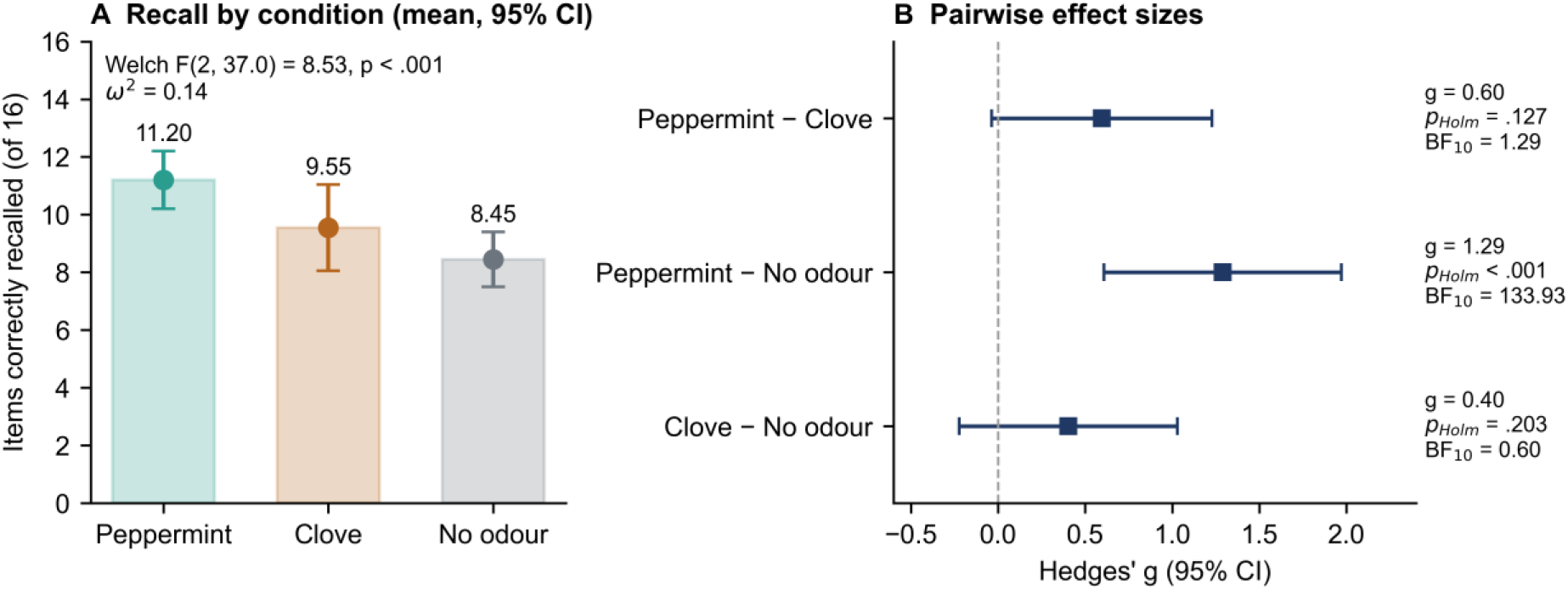
Recall by condition. (A) Mean number of items recalled with 95% confidence intervals. (B) Standardised pairwise differences (Hedges’ *g*) with 95% confidence intervals, Holm-adjusted *p* values and Bayes factors. The dashed line marks no difference.

Pairwise comparisons are summarised in Table 2 and Figure 2B. Participants exposed to peppermint recalled on average 2.75 more items than those exposed to no odour (95% CI [1.41, 4.09]; Welch *t*(37.9) = 4.16, *p*(Holm) < .001; *g* = 1.29, 95% CI [0.61, 1.97]), and the Bayes factor indicated very strong evidence for this difference (BF^10^ = 134). Recall after clove did not differ from no odour (difference = 1.10, 95% CI [-0.62, 2.82]; *p*(Holm) = .203; *g* = 0.40; BF^10^ = 0.60). The difference between peppermint and clove (1.65 items, 95% CI [-0.10, 3.40]) was of moderate size (*g* = 0.60) but not significant after correction (*p*(Holm) = .127; uncorrected *p* = .063), and the Bayes factor was inconclusive (BF^10^ = 1.29).

**Table 2.** Pairwise Comparisons Between Conditions.

| **Comparison** | **Δ M [95% CI]** | **Welch t (df)** | **p** | **p (Holm)** | **Hedges' g [95% CI]** | **BF<sub>10</sub>** |
|---|---|---|---|---|---|---|
| Peppermint vs. no odour | 2.75 [1.41, 4.09] | 4.16 (37.9) | < .001 | < .001 | 1.29 [0.61, 1.97] | 133.93 |
| Clove vs. no odour | 1.10 [-0.62, 2.82] | 1.30 (32.3) | .203 | .203 | 0.40 [-0.22, 1.03] | 0.60 |
| Peppermint vs. clove | 1.65 [-0.10, 3.40] | 1.92 (33.2) | .063 | .127 | 0.60 [-0.04, 1.23] | 1.29 |

*Note.* Welch-corrected independent-samples t tests; p (Holm) = Holm-adjusted p across the three comparisons; BF<sub>10</sub> = default JZS Bayes factor (Cauchy prior, r = .707).

## 4. Discussion

A single 30-s inhalation of peppermint essential oil was associated with markedly better immediate free recall of a 16-item visual array than no odour: participants recalled about 2.8 more items on average, a large standardised difference supported by very strong Bayesian evidence. Clove odour produced intermediate performance that did not differ reliably from either peppermint or no odour. These results are directionally consistent with previous reports that peppermint aroma enhances alertness and memory^12,13^ and that essential oils can have acute, odour-dependent effects on cognition^10,11,14^.

The pattern of pairwise results, however, limits the conclusions that can be drawn. If peppermint acted through a specific pharmacological or trigeminal mechanism, it should have outperformed clove as well as no odour. The peppermint–clove difference was of moderate size but inconclusive (BF^10^ ≈ 1), and clove itself showed a small, non-significant advantage over no odour. The data are therefore equally compatible with a general effect of odour exposure—greater arousal, novelty or engagement— graded by the intensity of stimulation, with peppermint’s menthol component providing additional trigeminal stimulation^17,18^. Within an adaptive-gain framework, moderate increases in noradrenergic arousal are expected to sharpen attention and enhance encoding^19^, consistent with the classic inverted-U relation between arousal and performance^20^.

Expectancy is a further, and important, alternative. Participants in the odour conditions inhaled from a bottle and knew they were receiving a scent, whereas the control group received nothing. Expectancies about essential oils can produce cognitive and physiological effects independently of the aroma^21,22^, and odours can act as primes whose influence depends on context and awareness^23^. Without a placebo odour and blinding, the present design cannot separate these routes. The neural plausibility of direct odour effects on memory is not in doubt^1,3,4^; what remains uncertain is whether brief exposure to a particular oil, rather than any salient odour experience, improves encoding of unrelated visual material.

Practically, these findings justify further work on low-cost olfactory interventions in learning environments, in line with evidence that olfactory enrichment can benefit memory over longer periods^15^ and that ambient odours influence mood and anxiety^24^. They do not yet justify recommendations for specific oils.

### 4.1. Limitations

First, the control group received no odour and no placebo bottle, and neither participants nor the experimenter were blind to condition; expectancy and experimenter effects cannot be ruled out. Second, testing took place in participants’ homes, so ambient odours, lighting and viewing distance were not standardised. Third, odour intensity, pleasantness and familiarity, as well as participants’ olfactory function, were not measured, and no physiological or self-report measure of arousal was obtained. Fourth, the sample was small; the study was powered only for large effects (*d* ≥ 0.91), so the non-significant clove comparisons should not be interpreted as evidence of no effect, and the large peppermint effect may overestimate the true effect size^28^. Fifth, a single array and a single immediate test were used, so effects on retention, recognition or other memory systems are unknown. Finally, participant-level data were not available for this re-analysis, which was therefore based on summary statistics; distributional checks and non-parametric confirmation could not be performed.

### 4.2. Conclusions

Brief peppermint odour exposure was associated with better immediate free recall of a visual array than no odour, but the study cannot attribute this benefit specifically to peppermint. A pre-registered laboratory replication with placebo and comparison odours matched for intensity and pleasantness, double-blind administration, olfactory screening and measures of arousal is the necessary next step.

## Declarations

### Ethics statement

Approved by the Non-Interventional Research Ethics Committee of Üsküdar University; the approval documentation is held by the authors and is available on request. Written informed consent was obtained from all participants.

### Data availability

Anonymised data are available from the corresponding author on reasonable request and will be deposited in a public repository upon acceptance.

### Author contributions (CRediT)

Bariş Akar: Conceptualization, Methodology, Investigation, Formal analysis, Visualization, Writing – original draft. Tuğba Altintaş: Methodology, Supervision, Writing – review & editing.

### Use of generative A

A generative AI tool was used to assist with re-analysis, figure preparation and language editing. The authors verified all analyses and text and take full responsibility for the manuscript.

